# The cumulative impact of passenger mutations on cancer development

**DOI:** 10.1101/2025.11.13.688369

**Authors:** Akshatha Nayak, Candace Chan, Ioannis Mouratidis, Ilias Georgakopoulos-Soares

## Abstract

The traditional binary classification of somatic mutations in cancer as either drivers or passengers overlooks the potential cumulative impact of smaller-effect mutations. Here, we analyze 2,263 whole-genome-sequenced primary tumors across 31 cancer types to assess the functional contribution of passenger mutations in cancer development. We find that in the absence of canonical driver mutations, passenger mutations in cancer genes are significantly enriched, exhibit higher predicted pathogenicity, and are associated with aberrant expression, splicing disruption, altered transcription factor binding, and clinical outcomes that resemble those in the presence of driver mutations. The accumulation of passenger mutations in tumor suppressor genes correlates with significantly reduced expression and poorer prognosis, mirroring the functional outcomes of driver mutations. Notably, this is a previously uncharacterized mechanism of tumor suppressor inactivation, in which the accumulation of somatic mutations results in its progressive inactivation. Our results support a continuum model of mutational impact, where the collective influence of passenger mutations contributes to oncogenesis and clinical outcomes. This work advocates for integrative cancer models that incorporate all somatic mutations to more accurately reflect the complexity of tumor evolution.

## Introduction

Cancer progression is characterized by the accumulation of somatic mutations ^1^. The prevailing dichotomy categorizes mutations as drivers, which directly contribute to cancer development, and passengers, which are considered incidental byproducts of the genomic instability inherent to cancer cells ^2^. Studies have identified a limited number of driver mutations that drive carcinogenesis per patient. In sarcomas, thyroid, and testicular cancers, there is approximately one driver mutation per patient, while in bladder, endometrial, and colorectal cancers, there are approximately four driver mutations per patient ^3^. Specifically, a study of 29 cancer types found that on average, only one to ten mutations are needed for a cancer to emerge. In contrast, passenger mutations can be hundreds or thousands of times more frequent, but their impact is supposedly negligible.

Emerging evidence suggests that the traditional driver-passenger mutation model oversimplifies the complexity of cancer development. Notably, up to 10% of cancer patients lack identifiable driver mutations, despite extensive efforts to catalog them across tens of thousands of tumors ^8^. Additionally, driver mutations are highly prevalent in healthy human tissues, and their prevalence increases with age ^4–6^. At older ages, a significant proportion of healthy cells across different tissues harbor driver mutations. Microdissections of tissues derived from aged people reveal that somatic cells with driver mutations proliferate faster and constitute a significant part of the tissue makeup, and that this phenomenon is widespread both across individuals and across tissues ^7^. Research has highlighted several possibilities to explain these phenomena and improve our understanding of cancer development, including the contribution of germline mutations, epigenetic alterations that promote tumorigenesis, and the role of the immune system in suppressing the expansion of potentially malignant cells ^9-11^. Previous cancer models have also examined weak driver mutations and deleterious passenger mutations on cancer progression and have found that mutational signature and subclonal architecture profiles can have aggregated effects in tumorigenesis ^8–10^.

Here, we tested the hypothesis that in certain cases, combinations of passenger mutations can collectively act in a driver-like manner. We investigated the distribution of passenger mutations near key cancer genes, comparing patient samples with and without driver mutations in them. Our analysis revealed an enrichment of passenger mutations, with higher predicted pathogenicity, in the absence of driver mutations, across cancer types. We report that the effect size of these differences is dependent on the cancer type and the cancer gene studied, and includes aberrant transcription factor binding, changes in gene expression, and splicing disruption. We also discovered that the accumulation of passenger mutations in tumor suppressor genes frequently results in their inactivation. For multiple cancer-related genes, the accumulation of passenger mutations results in overall survival outcomes that more closely resemble those of patients with driver mutations in these genes, compared to patients without either drivers or passengers. These findings reveal evidence of positive selection acting on passenger mutations in tumors lacking driver mutations. They support a model of cancer gene disruption as a continuum, challenging the prevailing binary classification of mutations (**Figure 1**).

**Figure 1:**
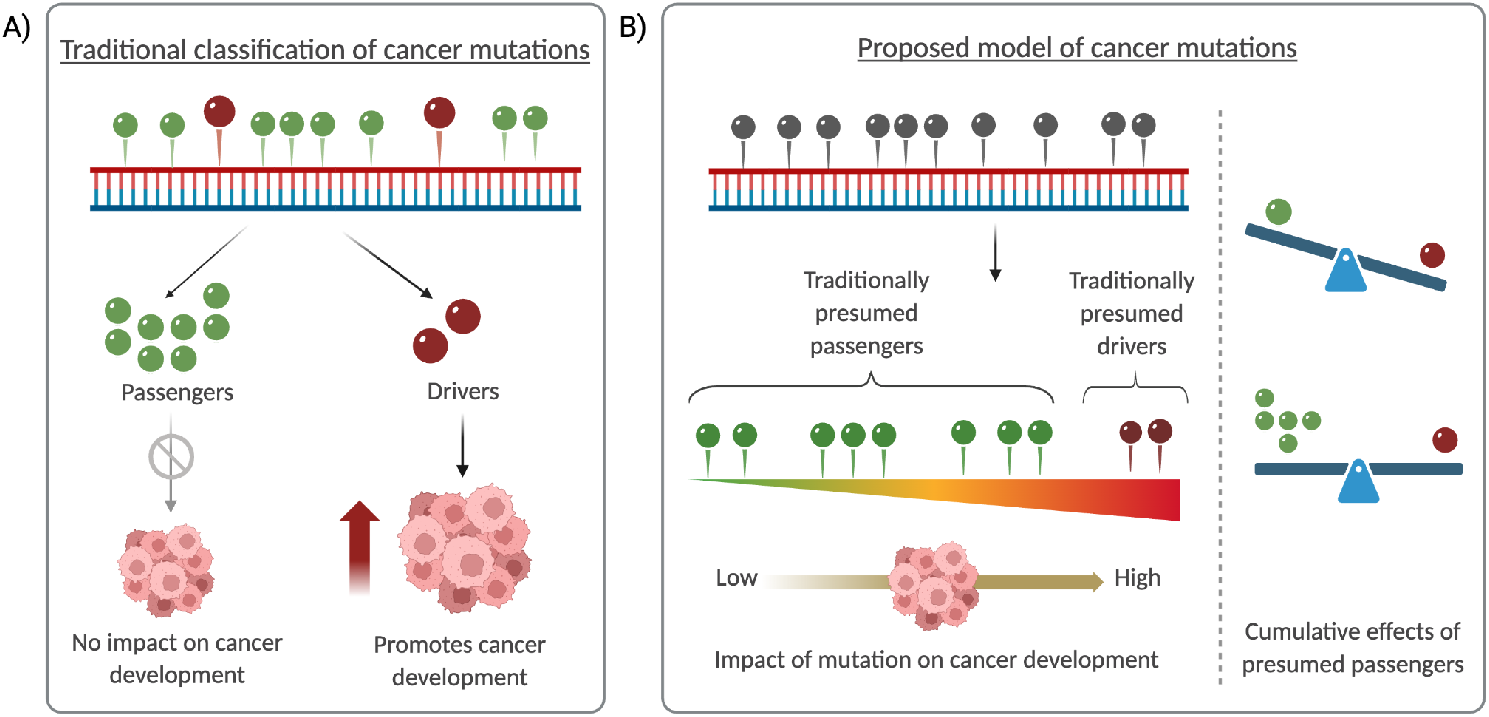
Incorporating a continuum of cumulative mutational impact in oncogenesis. **A)** Traditional binary classification of cancer mutations as drivers and passengers. **B)** Proposed model of cancer development with mutational effects on a spectrum.

## Results

We analyzed 2,263 whole-genome-sequenced (WGS) primary tumor samples across 31 different cancer types. We first annotated each primary tumor sample based on the driver mutations curated by the PCAWG consortium^11^. In accordance with previous findings ^3^, we observed substantial variation in the average number of driver mutations per primary tumor sample, with an overall mean of 5.3, but notably ranging from 11.3 in bladder cancers to 1.3 in thyroid cancers. Next, we were interested in examining whether, in the absence of driver mutations, we could identify evidence for smaller effect-size mutations that contributed to cancer development.

### Increased mutation rate in cancer genes when a patient lacks a driver mutation

Our hypothesis was that if passenger mutations play a driver-like role in the absence of a driver mutation, we would expect to find a higher passenger mutation burden in genes without a driver mutation than in genes with a driver mutation. To test this, we examined whether patients with a driver mutation in a specific cancer gene exhibit differences in the passenger mutation rate of that gene compared to patients without a driver mutation. We observed a statistically significant difference between the two groups, with patients who have driver mutations showing a significantly lower passenger mutation rate in that gene (Mann-Whitney U test, *p*-value<0.0001; **Supplementary Figure 1**). The results were unaltered when accounting for the genome-wide mutation rate differences between the two patient groups. When examining individual cancer tissues, we find that 12 out of 21 cancer tissues exhibited significantly more passenger mutations in the absence of drivers in cancer genes (Mann-Whitney U tests; **Figure 2a**). Notably, no tissue type demonstrated a significant enrichment of passenger mutations in the presence of driver mutations. Among the tissues with significant differences between the two groups, some of the largest enrichment in the mean adjusted passenger density was observed in esophageal (5.4-fold, *p*-value<0.0001) and head and neck (4.9-fold, *p*-value<0.0001) cancers.

**Figure 2:**
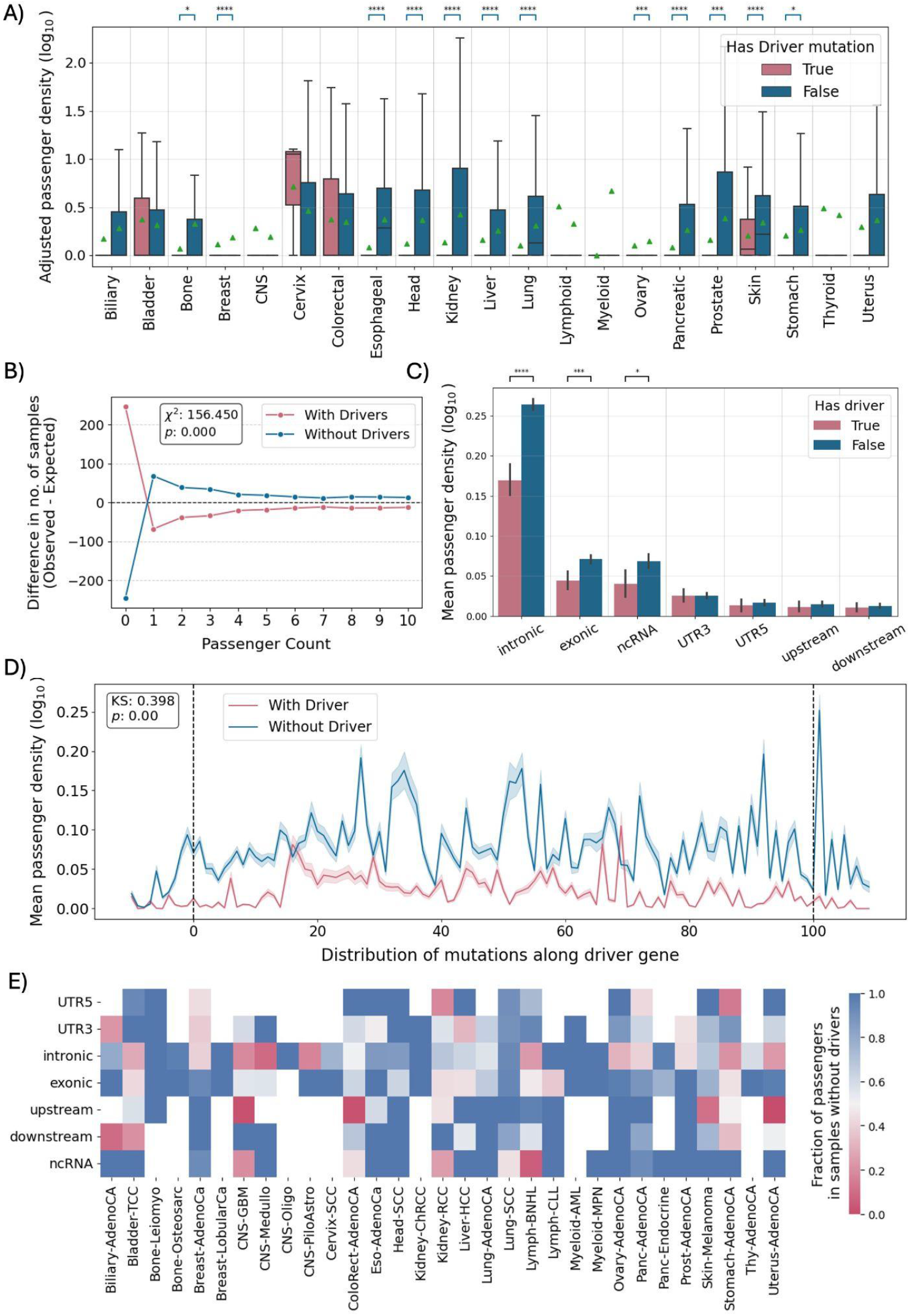
Passenger mutation density in samples with and without driver mutations. **(A)** Distribution of passenger mutation density for each cancer type. **(B)** Difference in observed and expected number of samples with varying passenger mutation counts in a cancer gene. **(C)** Mean passenger mutation density in different genic regions. **(D)** Passenger mutation density across 100 bins within the transcript, where 0 indicates the bin containing the TSS and 100 indicates the bin containing the TES. **(E)** Fraction of passenger mutation density in samples without driver mutations, measured as a ratio of average passenger density in samples without drivers to the sum of average passenger densities in groups of samples with and without drivers. Statistical significance is displayed as * for *p*-value<0.05, ** for *p*-value<0.01, *** for *p*-value<0.001, and **** for *p*-value<0.0001. The Mann-Whitney U test was used to estimate the statistical difference between groups.

To investigate the relative frequency of different passenger mutation burdens in samples lacking driver mutations compared to those with drivers, we stratified the samples based on the number of passenger mutations present in a cancer gene. We observed that samples without driver mutations in a cancer gene were less likely to exhibit an absence of passengers than samples with drivers. Consistent with the previous result (**Figure 2a)**, passenger mutations were more likely to be observed in samples without drivers in the cancer gene, compared to samples with drivers. While the difference in the observed and expected frequencies in this group diminished gradually with increasing numbers of passengers, the observation remained consistent across all non-zero passenger mutation count categories (Chi-square test of homogeneity, *p*-value=0.0; **Figure 2b**). These findings indicate that samples lacking driver mutations in cancer genes accumulate passenger mutations more often than expected, suggesting a compensatory mechanism for the absence of drivers.

To further examine whether the observed differences in passenger mutation rates between patients with and without driver mutations varied across distinct genic compartments, we stratified mutations by their genomic sub-compartments, including intronic, exonic, upstream, downstream, 5′ untranslated region (5′UTR), 3′ untranslated region (3′UTR), and non-coding RNA (ncRNA) regions. In all sub-compartments that we examined, we observed increased passenger mutation rates in the absence of driver mutations in a cancer gene (**Figure 2c**). Significant differences in passenger mutation rates were observed specifically within the intronic, exonic, and ncRNA regions (Mann-Whitney U tests, adjusted *p*-values<0.05; **Figure 2c**).

We also partitioned each gene into hundred equal-sized bins, and included 10 bins of the same size, upstream of the TSS and downstream of the TES. On examining the passenger mutational density in patient samples with and without driver mutations, we find that the elevated passenger mutation rate observed in the absence of a driver mutation in cancer genes is not localized to specific parts of cancer genes but spans the entire genic region and declines as we increase the distance upstream from the TSS and downstream from the TES (Kolmogorov-Smirnov test, *p*-value=0.0; **Figure 2d**). Notably, both upstream regions and regions downstream of transcription termination show increased passenger mutation density in the absence of a driver mutation. Additionally, when stratifying the analysis by cancer type, we observed consistent enrichment of passenger mutations in the absence of drivers across most cancer types and genic sub-compartments (**Figure 2e**), reinforcing the robustness and generality of this effect. These findings indicate that in the absence of a driver mutation, there are positive selective pressures for an increased number of passenger mutations, which likely partially disrupt cancer genes and contribute to the development of the cancer.

### Increased predicted pathogenicity of passenger mutations in the absence of drivers

We hypothesized that, in the absence of driver mutations, passenger mutations in these cancer genes might be under selection and would therefore exhibit larger pathogenicity scores than expected by chance, as measured by computational models. Therefore, in addition to examining differences in passenger mutation rates, we sought to determine whether the excess passenger mutations identified in cancer genes, occurring in the absence of driver mutations, have a larger effect size. Using Combined Annotation Dependent Depletion (CADD) scores, which are a computationally predicted measure of the deleteriousness of a variant, we find that the passenger mutations in cancer genes are predicted to be significantly more deleterious when a driver mutation is absent (Mann-Whitney U test, *p*-value<0.0001; **Figure 3a-b**). These findings are consistent across the cancer types examined, and in fourteen out of twenty-one cancer types, we find significantly higher deleteriousness for passenger mutations in the absence of a driver mutation than in its presence, indicating selection pressures even in the case of smaller effect sizes (Mann-Whitney U tests; **Figure 3c**). CNS cancers were the only type showing an opposite trend, which could be explained by the low mutation burden in this cancer type, as copy number variations (CNVs) and epigenetic alterations are thought to play a more prominent role in cancer progression ^12,13^ To further evaluate the cumulative deleterious effects of passenger mutations within each cancer gene, we used the sum of normalized CADD scores as a proxy measure. Consistent with observations based on individual variant-level pathogenicity, we found that passenger mutations exhibited a greater cumulative deleterious burden in the absence of driver mutations within the corresponding cancer gene (Mann-Whitney U test, *p*-value<0.0001) (**Figure 3d**).

**Figure 3:**
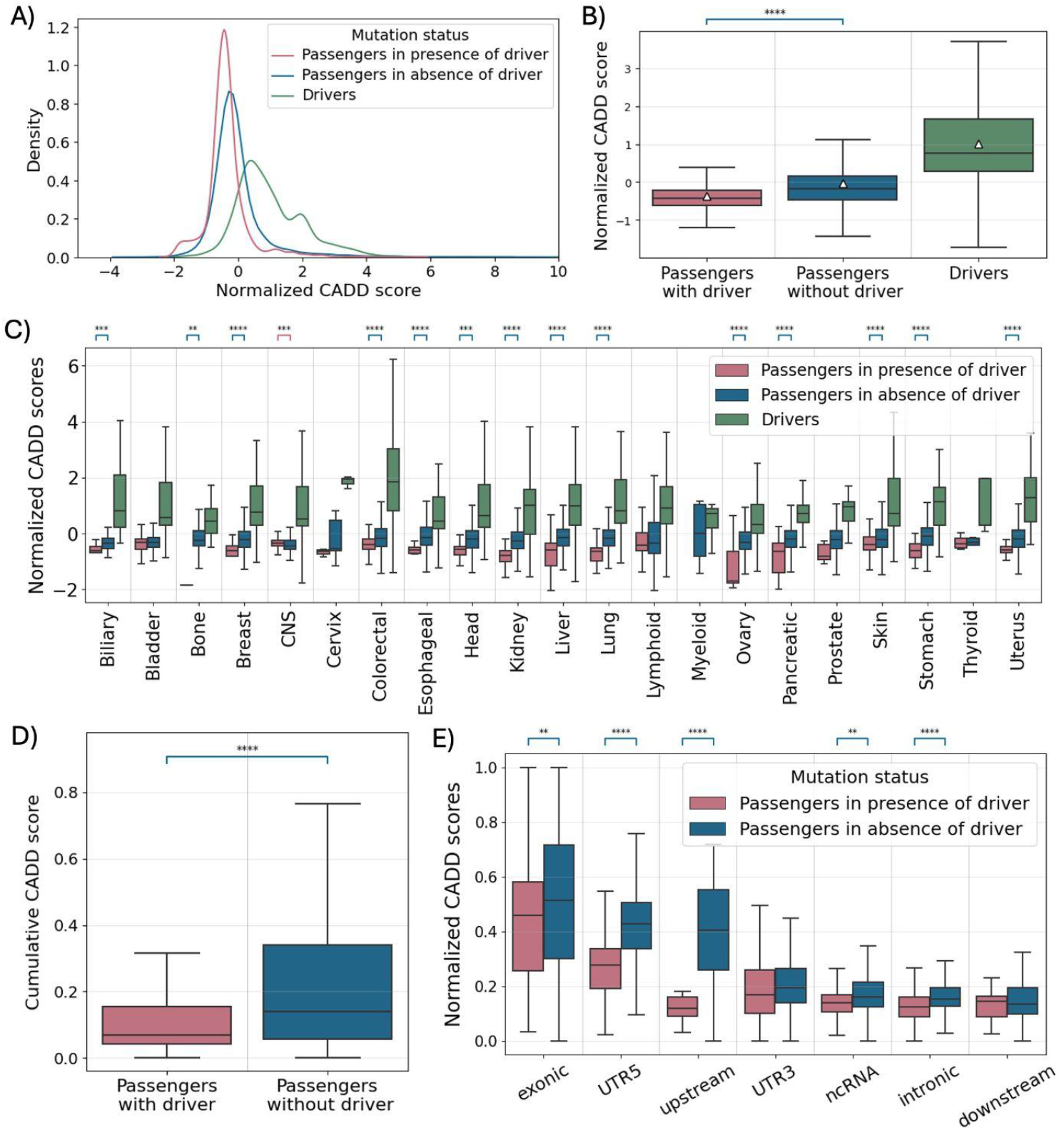
Pathogenicity profiles of passenger mutations in the presence or absence of driver mutations across cancer types. **(A)-(B)** Pan-cancer distribution of individual CADD scores for passenger mutations that emerge in the presence and absence of driver mutations. **(C)** Distribution of normalized CADD scores for passenger mutations in samples without driver mutation in a driver gene versus samples with driver mutation in the driver gene, by cancer tissue type. **(D)** Log-fold enrichment of median normalized CADD scores of passenger mutations in samples without driver mutation in a driver gene versus samples with driver mutation in the driver gene for each cancer subtype. **(E)** Distribution of normalized CADD scores for passenger mutations in samples without driver mutation in a driver gene versus samples with driver mutation in the driver gene, in different genic regions. Statistical significance is displayed as * for *p*-value<0.05, ** for *p*-value<0.01, *** for *p*-value<0.001, and **** for *p*-value<0.0001. The Mann-Whitney U test was used to estimate the statistical difference between groups.

To examine whether we could identify larger differences in the pathogenicity of passenger mutations in the absence of driver mutations, in specific genic compartments, we stratified the mutations by their genic sub-compartments, including intronic, exonic, upstream, downstream, 5′UTR, 3′UTR, and ncRNA regions. Interestingly, upstream regions and 5’UTR regions showed the strongest effect-size differences, while significant differences were also observed in ncRNA, exonic, and intronic regions (Mann-Whitney U tests; **Figure 3e**). We conclude that passenger mutations in cancer genes, in the absence of driver mutations, undergo selection pressures favoring increased pathogenicity. This observation underscores the functional relevance of integrating passenger mutations in cancer models, extending beyond the protein-coding regions into regulatory and non-coding compartments, thus highlighting previously underappreciated mechanisms contributing to tumor evolution.

### Promoter mutations affecting TF binding compensate for the absence of driver mutations

We also examined whether the primary tumor samples that lacked driver mutations in cancer genes were more likely to have promoter passenger mutations that disrupt TF binding. In our analyses, we excluded the annotated *TERT* promoter driver mutations, which have been characterized extensively and shown to be *bona fide* drivers. We used FABIAN-variant, a tool that can predict the effects of genetic variants on transcription factor binding ^14^. We focused our analysis on extended promoter upstream regions of cancer genes, which we defined as 1,000bp upstream of the TSS. We observe that for tumor samples that lack a driver mutation in the cancer gene, promoter passenger mutations cause significantly larger disruption in TF binding (Mann-Whitney U test, *p*-value<0.0001; **Figure 4a**). Differential analysis of individual transcription factors reveals that 107 out of the 1387 TFs exhibit significantly greater binding disruption by passenger mutations in the absence of driver mutations, whereas no TFs show significantly increased disruption by passengers in the presence of driver mutations (t-test, adjusted *p*-value<0.05; **Figure 4b**). We conclude that in the absence of driver mutations, partially compensatory mechanisms also include aberration of TF binding at cancer gene promoters.

**Figure 4:**
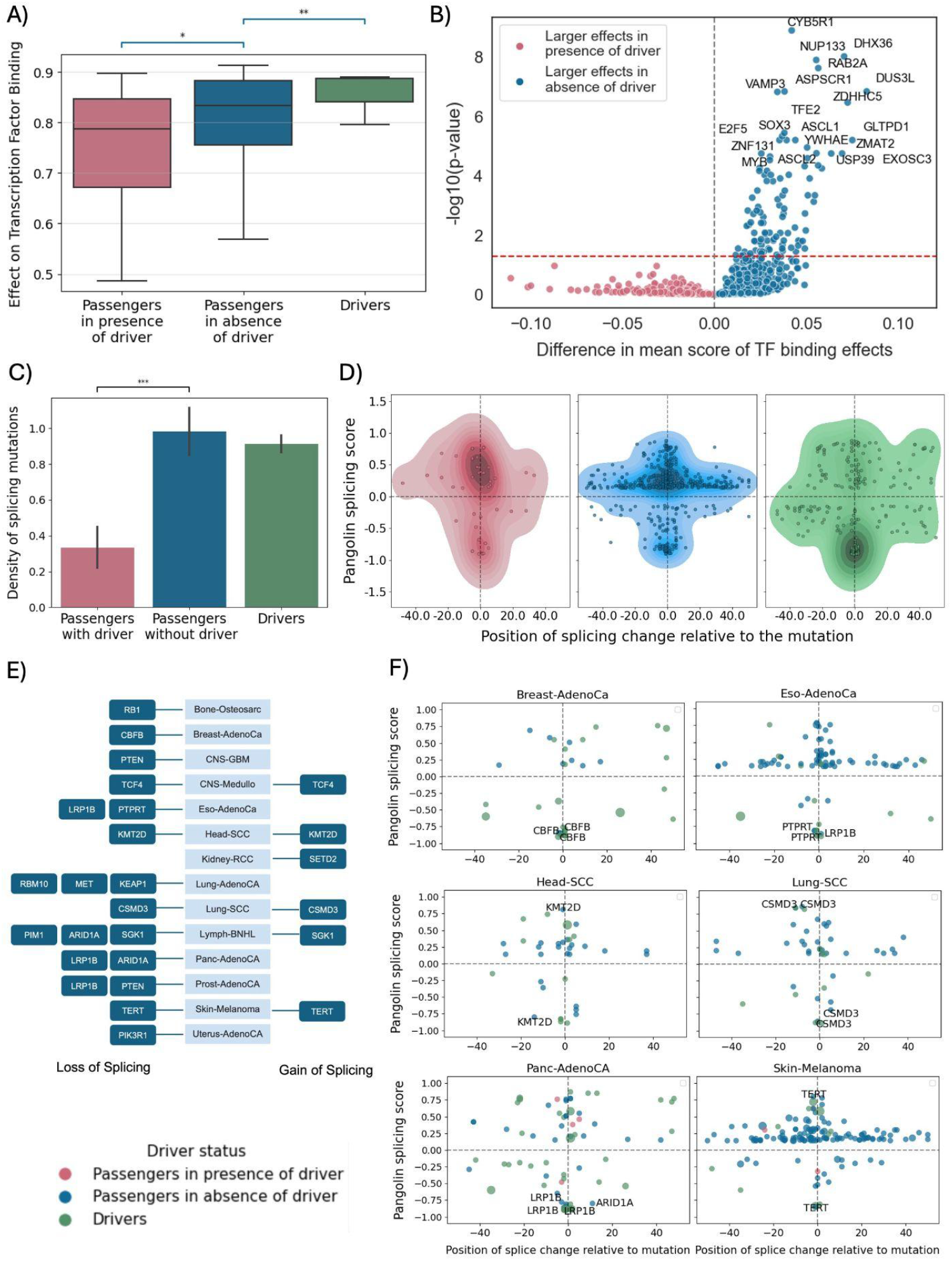
Excess of mutations causing splicing changes or altering transcription factor binding affinity in cancer genes lacking driver mutations. **A**. Effect of passenger mutations in the presence or absence of driver mutations, and of driver mutations, on transcription factor binding affinity. **B**. Differential effects on transcription factor binding affinity caused by passengers in the presence and absence of driver mutations. **C**. Density of passenger and driver mutations associated with splicing changes, with passengers stratified by the presence or absence of drivers. **D**. Effects of somatic mutations on splicing relative to the distance from the mutation, for individual cancer types. **E**. Genes which contain passenger mutations causing large splicing changes (absolute Pangolin score >= 0.8) in different cancer types. **F**. Distribution of large splicing changes (pangolin score >= 0.8) relative to the causative passenger and driver mutations for selected cancer types with high prevalence of splicing disruptive mutations. Statistical significance is displayed as * for *p*-value<0.05, ** for *p*-value<0.01, *** for *p*-value<0.001, and **** for *p*-value<0.0001.

#### Excess mutations impacting splicing in cancer genes in the absence of driver mutations

Next, we sought to determine whether the observed excess of pathogenic passenger mutations in cancer genes, in the absence of driver mutations, could induce aberrant splicing. Pangolin is a deep-learning-based method designed to predict changes in splice site strength due to genetic variants on RNA splicing ^15^. We leveraged Pangolin to gain insights into the consequences of these passenger mutations on splicing. In samples lacking a driver mutation in a given cancer gene, we observed a 2.9-fold increase in average frequency of passenger mutations predicted to induce splicing alterations, compared to samples with drivers (t-test, *p*-value<0.05; **Figure 4c-d**), suggesting positive selection for changes that lead to alternative splicing during cancer development. Although these mutations were associated with both loss and gain in splicing (**Figure 4d-f**), a majority of the passenger mutations were found to cause splicing gain (**Figure 4d**). Additionally, we identified several genes across different cancer types, in which we find passenger mutations that cause large splicing disruptions (absolute Pangolin score >= 0.8) (**Figure 4e, Supplementary Figure 2**). Notably, *LRP1B, PTEN*, and *ARID1* genes harbored passengers causing large splicing disruptions in more than one cancer type (**Figure 4e**). Our results indicate pervasive splice site changes in the absence of driver mutations, advocating for the need of holistic models that integrate passenger mutations into them.

### Incorporation of passenger mutations in cancer genes enables improved prognosis

To investigate the potential prognostic value of passenger mutations, we compared the median survival time (based on overall survival as the endpoint) of patients with no mutations in a cancer gene, patients having only passenger mutations but no driver mutations, and patients with driver mutations in the gene. Our analysis was limited to a subset of 710 patients due to the limited availability of their corresponding clinical data. We find that patients with only passenger mutations in a cancer gene had significantly lower survival times than patients without any mutations in the gene (Mann-Whitney U test, *p*-value < 0.0001; **Figure 5a**). Further analysis using the Kaplan-Meier estimator revealed significant differences in overall survival outcomes across mutational categories. As expected, patients with a driver mutation in a cancer gene exhibited worse survival outcomes than patients with no mutations in the corresponding gene (log-rank test, *p*_2_=0.0). However, patients with passenger mutations in the absence of driver mutations also demonstrated significantly worse survival outcomes (log-rank test, *p*_1_=0.0; **Figure 5b)**, suggesting that passenger mutations may independently carry prognostic relevance.

**Figure 5:**
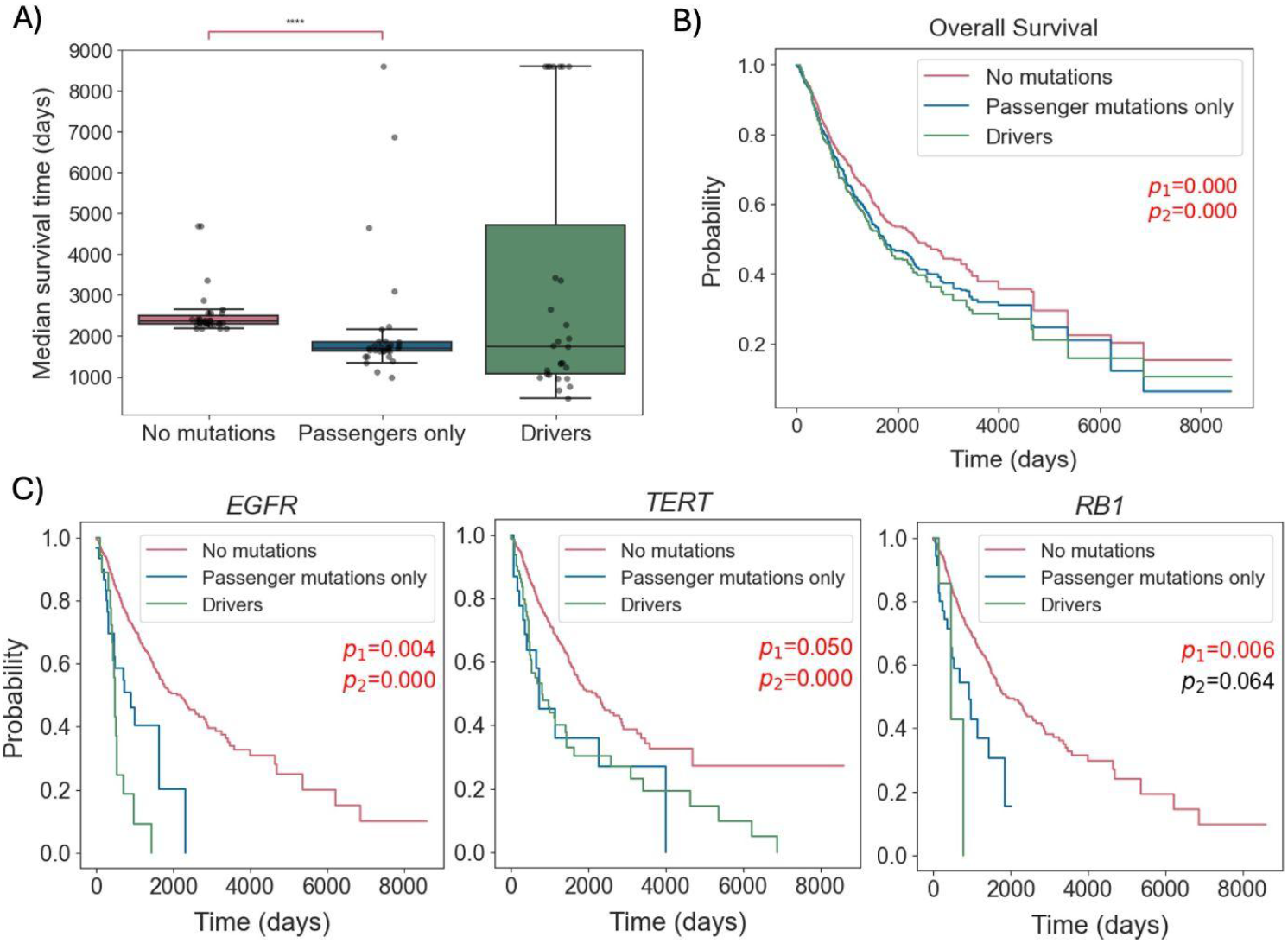
Effect of passenger mutations on overall survival (OS) **(A)** Distribution of median survival time (in days) of patients. Each data point corresponds to the median survival for a particular cancer gene. **(B)** Kaplan-Meier Survival analysis **(C)** Kaplan-Meier Survival analysis for genes where passenger mutations lead to a statistically significant decrease in the overall survival (OS) of the cancer patient. Patients are grouped into 3 categories: patients without any mutations, patients with passenger mutations only, and patients with driver mutations. *p*_1_ refers to the *p*-value for groups “No mutation” vs “Passenger mutations only”, and *p*_2_ refers to the *p*-value for groups “No mutation” vs “Drivers”. Statistical significance is displayed as * for *p*-value<0.05, ** for *p*-value<0.01, *** for *p*-value<0.001, and **** *for p*-value<0.0001.

We further examined the clinical utility of passenger mutations in certain key cancer genes. Similar to the aggregated analysis, we categorized patients into three groups. After accounting for multiple testing, we identified three genes, namely *EGFR, TERT*, and *RB1*, in which patients with only passenger mutations exhibited significantly poorer survival outcomes compared to those without any mutations (Log-rank tests, *p*-values < 0.05; **Figure 5c**). *EGFR* is a receptor tyrosine kinase that belongs to the ErbB family of proteins. Patients with disruption of EGFR often benefit from EGFR Tyrosine Kinase Inhibitors, which significantly improve progression-free survival compared to chemotherapy ^16^. *RB1*, the retinoblastoma tumor suppressor gene, is a key regulator of cell-cycle processes. Accumulating evidence indicates that RB1 status is a critical determinant of cellular sensitivity to diverse antitumoral therapies, such as kinase inhibitors and immunotherapy ^17^. Finally, TERT promoter (TERTp) mutation status has been shown to significantly impact cancer development. However, aberrant telomerase activity is observed in several cancer types, even in the absence of frequent TERTp driver mutations, suggesting alternative or epigenetic mechanisms likely contributing to tumorigenesis ^18^. The passenger mutations within the TERT gene may provide additional insights into these mechanisms and help understand the molecular basis of telomerase activation in cancers lacking canonical drivers. Our findings thus highlight the prognostic significance of passenger mutations in cancer genes.

### Clusters of passenger mutations inactivate tumor suppressor genes

We hypothesize that passenger mutations are positively selected during cancer development, due to cumulative effects that disrupt gene expression. We used RNA-seq data from the primary tumor samples and investigated whether the number of passenger mutations in these genes impacted the expression levels of oncogenes and tumor suppressors when no driver mutations were present. We find a distinct trend, with the number of passenger mutations occurring in tumor suppressor genes being strongly correlated with lower expression levels (Kruskal-Wallis test, p-value=0; **Figure 6a**). Importantly, the accumulation of passenger mutations results in the inactivation of the tumor suppressors. For instance, we observe a 2.5-fold reduction in tumor suppressor gene expression in the presence of 3-5 passenger mutations, and an approximately 5-fold reduction when there are 5-10 passenger mutations present, relative to genes without any mutations, corresponding to a level of transcriptional silencing consistent with functional inactivation. We also observe a weaker association between the number of passenger mutations occurring in an oncogene and increased expression (Kruskal-Wallis test, p-value=0.04; **Figure 6a**). Notably, this effect is driven purely by the number of mutations and not by specific passenger mutations recurring across patient samples, indicating that the accumulation of a multitude of different passenger mutations with small individual effects can disrupt gene function. We conclude that passenger mutations contribute to tumor suppressor inactivation and oncogene upregulation in a dosage-dependent manner.

**Figure 6:**
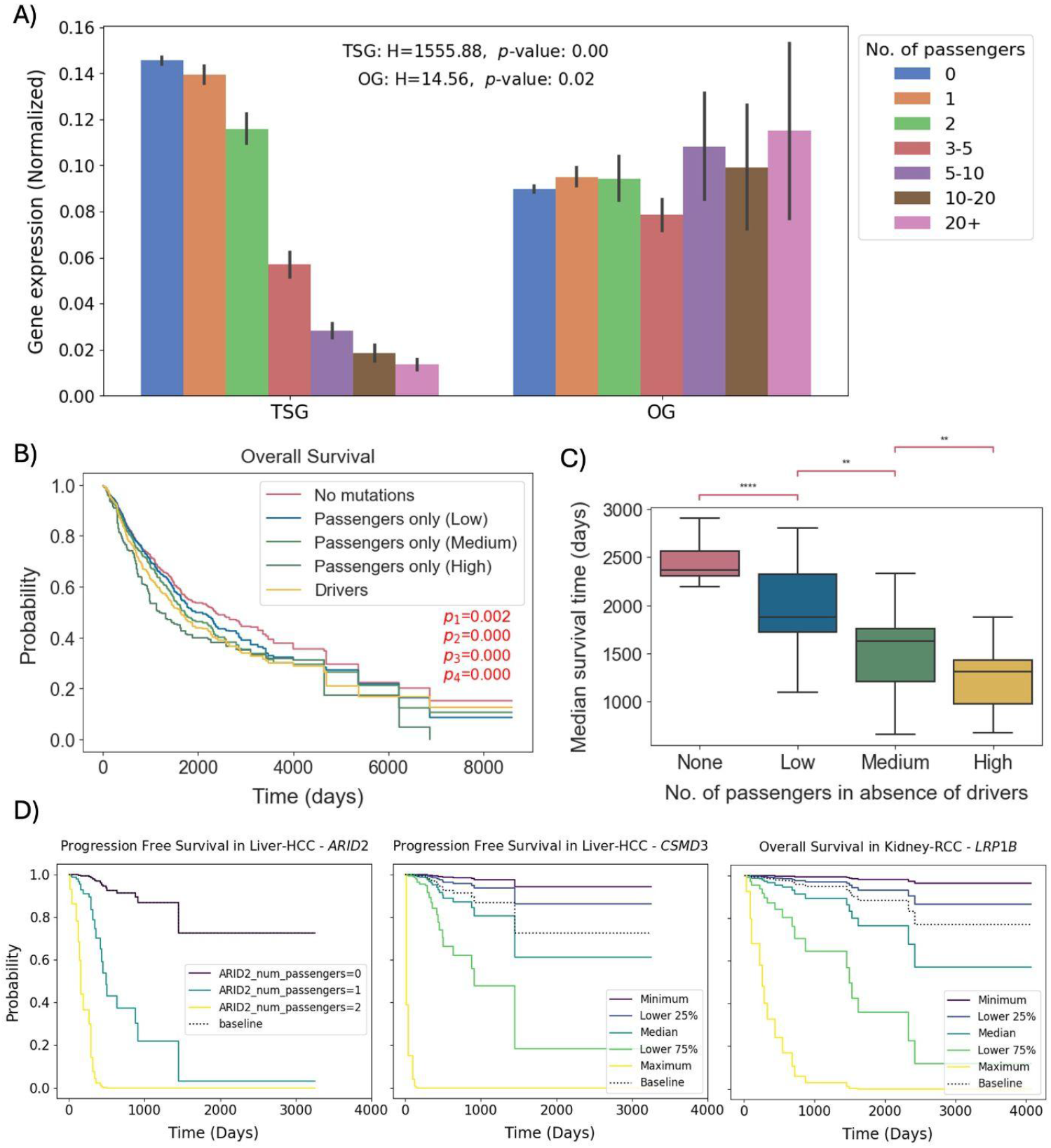
Cumulative effects of passenger mutations. **(A)** Relationship between the number of passenger mutations in the absence of drivers and gene expression levels in cancer genes. Results shown for tumor suppressor genes (TSG) and oncogenes (OG). Statistical significance was estimated using the Kruskal-Wallis test. **(B)** Kaplan-Meier survival analysis stratified by the number of passenger mutations in the absence of drivers. *p*_1_ refers to the *p*-value for groups “No mutation” vs “Passengers only (Low)”, *p*_2_ refers to the *p*-value for groups “No mutation” vs “Passengers only (Medium)”, *p*_3_ refers to the *p*-value for groups “No mutation” vs “Passengers only (High)”, and *p*_4_ refers to the *p*-value for groups “No mutation” vs “Drivers”. **(C)** Distribution of median survival time (in days), stratified by the number of passenger mutations in the absence of drivers. Statistical significance is displayed as * for *p*-value<0.05, ** for *p*-value<0.01, *** for *p*-value<0.001, and **** for *p*-value<0.0001. **(D)** Partial effects of the number of passengers as a covariate in the Cox proportional hazards model for progression-free survival and overall survival.

### Accumulated passenger mutations in cancer genes adversely impact clinical outcomes

Next, we examined whether the accumulation of passenger mutations in cancer genes is associated with clinical outcomes of cancer patients. We grouped patients either by the presence of a driver mutation in the cancer gene or, in the absence of a driver mutation, we stratified patients into quartiles based on the number of passenger mutations in the same gene. These quartiles were consolidated into low (Q1), intermediate (Q2-Q3), and high (Q4) passenger-mutation groups (**Figure 6b-c**). We found that in patients without driver mutations in a cancer gene, survival outcomes became, on average, progressively worse with an increasing number of passenger mutations in the gene (log-rank tests, *p*-values < 0.005, **Figure 6b;** Mann-Whitney U tests, *p*-values < 0.01 and Jonckheere-Terpstra test, *p*-value = 0.0, **Figure 6c**). To investigate the independent contribution of the number of passenger mutations in a cancer gene on the survival outcomes, adjusting for other genetic, clinical, and demographic covariates, we fitted a Cox proportional hazards model. After multiple testing hypothesis correction, the number of passenger mutations in the *ARID2* and *CSMD3* genes in hepatocellular carcinoma patients, as well as in the *LRP1B* gene in renal cell carcinoma patients, emerged as statistically significant covariates associated with poorer prognosis (adjusted *p*-value < 0.05; **Figure 6d**). Notably, in *ARID2*, the presence of even a single passenger mutation, found in different genomic loci across the gene, was found to be sufficient to reduce the survival probability of hepatocellular carcinoma patients below the baseline hazard, whereas in *CSMD3* and *LRP1B* genes, a moderately higher number of passenger mutations, ranging between the 25th and 50th percentiles, was required for a decrease in survival probability below the baseline hazard. Collectively, these findings suggest that passenger mutations in cancer genes are an independent and clinically relevant determinant of survival outcome, highlighting their potential utility in patient risk stratification.

## Discussion

The dichotomy model of driver and passenger mutations primarily focuses on non-synonymous mutations in cancer genes, copy number aberrations, and structural variants as the drivers of cancer development ^19–21^, while the remaining mutations are considered bystanders. In this study, we provide evidence that subsets of passenger mutations, traditionally considered biologically neutral or even detrimental to tumor growth, are subject to positive selection and can exert functional consequences during cancer development. By analyzing 2,263 WGS tumors, we demonstrate that in the absence of known driver mutations, cancer genes harbor a significantly higher burden of passenger mutations. These mutations are enriched across genic regions and regulatory elements and exhibit elevated predicted pathogenicity. This trend is consistent across multiple cancer types, suggesting that passenger mutations are under selective pressure and may play a compensatory role when driver mutations are absent. These findings challenge the canonical driver-passenger dichotomy and suggest that passenger mutations can collectively contribute to oncogenesis in a context-dependent manner.

Our findings suggest that smaller, additive effects can collectively disrupt cancer genes and contribute to tumorigenesis. Tumor suppressor genes typically require biallelic inactivation ^22^, and thus the accumulation of passenger mutations may contribute to gene disruption of both alleles in accordance with the two-hit hypothesis. In contrast, oncogenes are typically activated by a single mutant allele ^23^ and are therefore less vulnerable to additional disruptions; nonetheless, we still observe a weaker but significant association with the accumulation of passenger mutations. Our work also shows that the disruption of these genes by the accumulation of passenger mutations can be multifactorial, and these mutations are dispersed throughout the genomic sub-compartments and regulatory elements. Current models of tumor suppressor gene inactivation do not account for the collective effects, and their incorporation in future models can provide clinical benefits in diagnosis, prognosis, patient stratification, and treatment choice.

We have shown that smaller-effect somatic mutations are significantly more likely to cause aberrant transcription factor binding and splicing, and their accumulation can disrupt tumor suppressor genes. Consistent with our findings on splicing, prior research has demonstrated that synonymous mutations can serve as oncogenic drivers ^24,25^. Notably, we identify 137 transcription factors with significantly disrupted binding sites in the promoters of cancer genes lacking driver mutations, whereas no significant disruption of transcription factor binding is observed in cancer genes with driver mutations, suggesting a pervasive, compensatory mechanism involving smaller regulatory effects. Studies have previously shown that perturbation of transcription factor binding sites, such as ETS binding sites, can have oncogenic potential and associations with patient survival ^26,27^, consistent with our findings.

Additionally, we find that the incorporation of the collective passenger mutation burden of cancer genes in clinical models can better inform the survival of cancer patients. Rather than a binary model of cancer gene disruption, there exists a continuum of functional impact. This adds to previously proposed models such as polygenic mini-driver models, which concentrate on finding additional cancer genes and smaller effect sizes ^8,28^, but differs in that our study focuses on the importance of cumulative mutational effects in them. Furthermore, while deleterious effects and improved clinical outcomes in tumors with accumulated passenger mutations have been characterized ^29^, our work suggests advantageous effects for tumor development and worse clinical outcomes when the passenger mutations accumulate specifically in cancer genes ^29^.

In this work, we emphasize a shift from a discrete to a continuum-based model, where the distinction between driver and passenger is, in certain cases, not binary but context-dependent. Future work will focus on developing a holistic model of cancer development that integrates all mutations by quantifying their weighted, collective impact on tumor behavior. These models will account for the cumulative effects of low-impact mutations, context-specific dependencies, and the interplay between coding, non-coding, structural, and epigenetic alterations to better capture the complexity of cancer evolution.

## Methods

### Datasets used

Somatic mutations from 2,778 primary tumor samples across 21 tissues were analyzed using the hg19 reference human genome. Substitution and indel mutations, as well as CNVs, were analyzed from Pan-Cancer Analysis of Whole Genomes (PCAWG) from ICGC and TCGA. Clinical data were derived from the ICGC data portal, and survival data were collected from the TCGA Pan-Cancer Clinical Data Resource (TCGA-CDR) ^30^.

Driver genes for each of the PCAWG cancer cohorts were obtained from the IntoGen database ^2^. Driver genes that were mutated in at least 2% of the samples were selected. Genes that were mutated in at least 2% of samples within a given cohort were retained for further analysis. Subsequently, the top 10 most frequently mutated driver genes were selected for each cohort. In cases where multiple genes shared the same mutation frequency as the 10th-ranked gene, all such genes were included. Although the TERT gene is a well-established driver in several cancer types, it was not listed in the IntoGen dataset. Therefore, TERT was manually added to the driver gene lists for the following cancer types: Bladder-TCC, CNS-GBM, CNS-Medullo, Head-SCC, Liver-HCC, Skin-Melanoma, and Thy-AdenoCA, in accordance with its classification as a driver gene for these cancer types by the PCAWG consortium. For pancancer analyses, all driver genes mutated in at least 2% of the patients across the pancancer cohort were selected, resulting in a set of 15 driver genes.

Driver mutations for each patient were obtained from the ICGC and TCGA datasets. Genic regions were derived from RefSeq ^31^. Genic regions annotation of somatic mutations was performed using Annovar ^32^ (latest version as of Dec 5, 2024), and their predicted pathogenicity was assessed using scores from the CADD database ^33^ (CADD v1.7 for GRCh37/hg19).

### Mutational burden in cancer genes with and without driver mutations

To compare mutational burden per cancer gene, mutations were divided into two groups: samples containing at least one driver mutation in the gene, and samples without any identified driver mutations in the same gene, as annotated in the PCAWG dataset. Mutational density was calculated separately for each group, adjusting for genome-wide differences in mutation rates. Each data point represents the mutation density of the gene in an individual patient sample within that group. Differences in mutational burden between groups were then assessed using Mann-Whitney U tests to evaluate the statistical significance based on the driver mutation status. The same analysis was repeated in individual genomic subcompartments, namely 5’UTRs, 3’UTRs, intronic, exonic, upstream (1000 bp), and downstream (1000 bp) genic regions, and in ncRNAs. The Benjamini-Hochberg procedure was used for multiple testing correction.

Let,

*n*= No. of passenger mutations in the gene

*L*= Length of the Genome

*N* = Total no. of mutations across the genome

*l* = Gene length

*C* = CNA burden, i.e., fraction of genome length having copy number alterations

*M* = Mutation density adjusted for genome-wide mutation rate and CNA burden

Mutation rate 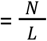

Expected rate in gene = Mutation rate * Gene length 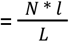

Mutation density after accounting for the expected mutation rate in the gene 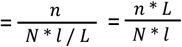

Mutation density after adjusting for CNA burden, 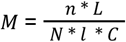

### Frequency distribution of passenger mutation counts

For the same groups of samples as described previously, we tabulated the observed frequencies of samples with varying numbers of passenger mutations (0-10). A Chi-square test of independence was then performed to calculate the expected frequencies under the null hypothesis of independence. The resulting difference in observed and expected frequencies for the two groups was visualized, and the chi-square statistic, along with the *p*-value, was reported to assess the statistical significance.

### Passenger density relative to the Transcription start (TSS) and end (TES) sites

Each cancer gene was divided into 100 equal-sized bins to correct for differences in gene length, to which 10 additional bins of equal length were added upstream and downstream of the TSS and TES, respectively. To estimate the 95% confidence interval for the mean, bootstrapping was performed with 100 iterations. For each bin, the adjusted passenger mutation density for each sample was estimated by adjusting for the gene length (*l*), genome-wide mutation rates (*N*), and copy number alteration burden (*C*). The Kolmogorov-Smirnov (KS) test was used to evaluate the statistical difference between the two distributions.

The adjusted mutation density in the i^th^ bin is, 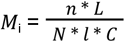 where, *n*= No. of passenger mutations in the i^th^ bin of the gene, and other variables are as defined previously.

### Predicted mutation pathogenicity in the presence or absence of driver mutations

The pathogenicity of mutations in each of these three groups was quantified using Combined Annotation Dependent Depletion (CADD) scores ^33^. For each cancer type and patient, somatic mutations in cancer-related genes were stratified into three groups: (1) mutations annotated as drivers according to the PCAWG dataset, (2) passenger mutations occurring in cancer genes co-occurring with driver mutations but not themselves annotated as drivers), and (3) passenger mutations occurring in cancer genes without any identifiable driver mutations co-occurring. To facilitate comparative analysis, CADD scores were z-score normalized within each gene. For cumulative analysis, min-max normalization was applied to constrain the values to a positive range, to improve interpretability in additive contexts. The analysis was also repeated for different cancer types and in different genomic sub-compartments. In all cases, statistical significance was estimated using Mann-Whitney U tests, and the Benjamini-Hochberg procedure was used for multiple testing correction.

### Characterization of somatic mutations in the vicinity of splice sites

The Pangolin software (v1.0.1) was used for predicting splicing changes caused by each somatic mutation ^15^. The software provides the locations of predicted splicing change relative to each mutation, and a score corresponding to the magnitude and direction (loss or gain) of the splicing change. The absolute value of the Pangolin splicing scores was obtained as recommended by the authors at the vicinity (50 base pairs on either side) of (1) driver mutations in a cancer gene, (2) mutations co-occurring with a driver mutation in the same cancer gene in that sample, and (3) mutations in cancer genes without co-occurring driver mutations. Similarly, a False Sign Rate (FSR) of 5% was selected as the cutoff, resulting in sites with an absolute splicing score of 0.14 and above being considered splicing sites. The density of splicing changes caused by mutations in different groups based on this cut-off was compared, and the statistical significance was estimated using the t-test.

The adjusted density of splicing changes in a gene, per gigabase (*S*) is calculated as,

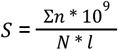

where *n* is the number of splicing changes caused by each mutation, and Σ*n* corresponds to the sum of all splicing changes caused by all the mutations in a gene. This was adjusted for the gene length (*l*), genome-wide mutation rates (*N*).

### Predicted effects of somatic mutations in promoter regions on transcription factor binding

FABIAN-Variant (web service) was used for predicting variant effects on transcription factor binding affinity ^14^. The web-based tool predicts the impact of a variant on the binding affinity of 1,387 human transcription factors. Similar to the splice site analysis, we used groups of patients with (1) driver mutations in a cancer gene, (2) mutations co-occurring with a driver mutation in the same cancer gene in that sample, and (3) mutations in cancer genes without co-occurring driver mutations. For this analysis, we prioritized mutations within the 1,000 bases upstream of the transcription start sites (TSS). To assess the potential impact of these mutations on transcription factor (TF) binding, we defined the effect size as the maximum predicted disruption caused by a mutation across a panel of 1,387 TFs. We then examined these effects in each of the three groups.

For each transcription factor, we compared the mean absolute effect of passenger mutations occurring in the presence and absence of driver mutations in the gene, across the pancancer cohort, using the t-test. This analysis was conducted for all 1,387 transcription factors, followed by multiple testing correction using the Benjamini-Hochberg method, to derive the adjusted *p*-values. To visualize the results, we generated a volcano plot, with the difference in mean absolute transcription factor binding effects represented on the x-axis and the negative logarithm of the adjusted p-values on the y-axis. The 20 transcription factors exhibiting the most statistically significant differential disruption were annotated on the plot.

### Overall survival analysis

Patients were classified into three groups: (1) patient samples with driver mutations in the studied cancer gene, (2) patient samples with passenger mutations but no driver mutations in the studied gene, and (3) patient samples without any mutations in that gene. A subset of 710 patients was used for the analysis due to the limited availability of the corresponding clinical data. The median survival time was calculated for patients within each of the three groups for every gene, and the overall distribution across all genes was analyzed. For patients who did not experience the event (death) during the study period, survival time was censored at the study’s end. Statistical significance was estimated using the Mann-Whitney U test.

Kaplan-Meier survival analysis curves for each gene were generated using the *lifelines* library in Python for the three groups. For this analysis, we shortlisted cancer genes that had at least 5 samples in group 1 and at least 10 samples in groups 2 and 3. An integrated analysis was also conducted using the set of driver genes for the pan-cancer cohort. Statistical significance was estimated using the log-rank test to compute *p*-values, adjusting for multiple testing using the Bonferroni correction.

### Effect of the number of passenger mutations on gene expression and survival outcomes

Gene expression analysis was performed for cancer genes at the pancancer level. Expression data was obtained from the *tophat_star_fpkm*.*v2_aliquot_gl*.*tsv*.*gz* file in the PCAWG dataset, which provides FPKM-normalized expression values representing the average of TopHat2 and STAR-based RNA-seq alignments. To enable combined analysis across genes, gene expression values were further normalized using min-max scaling on a per-gene basis. For each gene, patients without driver mutations in the gene were selected and stratified by the number of passenger mutations. The mean normalized expression was measured for tumor suppressor genes and oncogenes separately. Statistical significance of the differences in gene expression associated with passenger mutation burden was estimated using the Kruskal-Wallis test.

Similar to the previous survival analysis, patients were categorized into three primary groups: (1) those with no mutations, (2) those harboring only passenger mutations, and (3) those with driver mutations in cancer genes. To evaluate the effect of the number of passenger mutations on overall survival, we further stratified group 2 by low, medium, and high number of passengers, resulting in a total of 5 groups. This stratification was determined according to the quartile distribution of passenger mutation counts, with the first quartile designated as low, the second and third quartiles as medium, and the fourth quartile as high. Kaplan-Meier survival analysis curves were generated using the *lifelines* library in Python for the five groups, and statistical significance was estimated using the log-rank test to compute *p*-values.

We further calculated the median survival time for patients within each of the five groups for every gene and analyzed the overall distribution across all genes. For patients who did not experience the event (death) during the study period, survival times were right-censored at the study endpoint. Statistical significance of pairwise differences in survival distributions was assessed using the Mann-Whitney U test. To evaluate the presence of a monotonic trend in survival with increasing passenger mutation burden, we applied the Jonckheere-Terpstra trend test as implemented in the *clinfun* package in R.

### Proportional hazard associated with the number of passenger mutations

Hazard modelling was performed using the lifelines^34^ library in Python for each cancer type, using progression-free interval (PFI) and overall survival (OS) as clinical endpoints or events. We applied a Cox proportional hazards model to assess the association between survival and multiple covariates, including the presence of driver mutations in individual cancer genes, the number of passenger mutations per gene, event occurrence, time to event, age at diagnosis, sex, the total number of driver mutations across the genome, the total mutational burden, and copy number alteration (CNA) burden. Genes with passenger mutations in at least ten samples were included in the model. Genes for which the number of passenger mutations was identified as a statistically significant covariate (*p* < 0.05), after adjusting for multiple testing using the Bonferroni correction, were selected for further analysis of their partial effects on the outcome. For genes exhibiting a wide range in the number of passenger mutations, the partial effects on survival were evaluated at the 0^th^, 25^th^, 50^th^, 75^th^, and 100^th^ percentiles of passenger mutation counts.

## Code Availability

The GitHub code and all the related material for this work is provided at: https://github.com/Georgakopoulos-Soares-lab/cancer_model

## Author Contributions

A.N., I.M., and I.G.S. jointly conceived the study. A.N. performed data curation, formal analysis, and interpreted the results. A.N., C.C., and I.G.S. designed the methodology. A.N. generated the code, statistical analyses, and visualizations. A.N. and I.G.S. wrote the manuscript with input from I.M. and C.C. I.G.S. acquired funding for the project and provided supervision.

## Acknowledgements

This work utilized data generated by the Pan-Cancer Analysis of Whole Genomes (PCAWG) project ^11^. The results published here are based upon data generated by the TCGA Research Network and accessed through dbGaP (phs000178.v11.p8).

