## Supplementary Figures for "The cumulative impact of passenger mutations on cancer development"

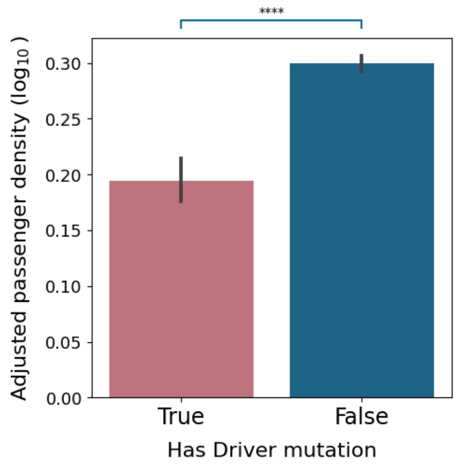


**Supplementary Figure 1: Aggregate passenger mutation density between groups of samples with and without driver mutation in a cancer gene.** Mann-Whitney U tests with p-value is displayed as **** for p-value<0.0001.

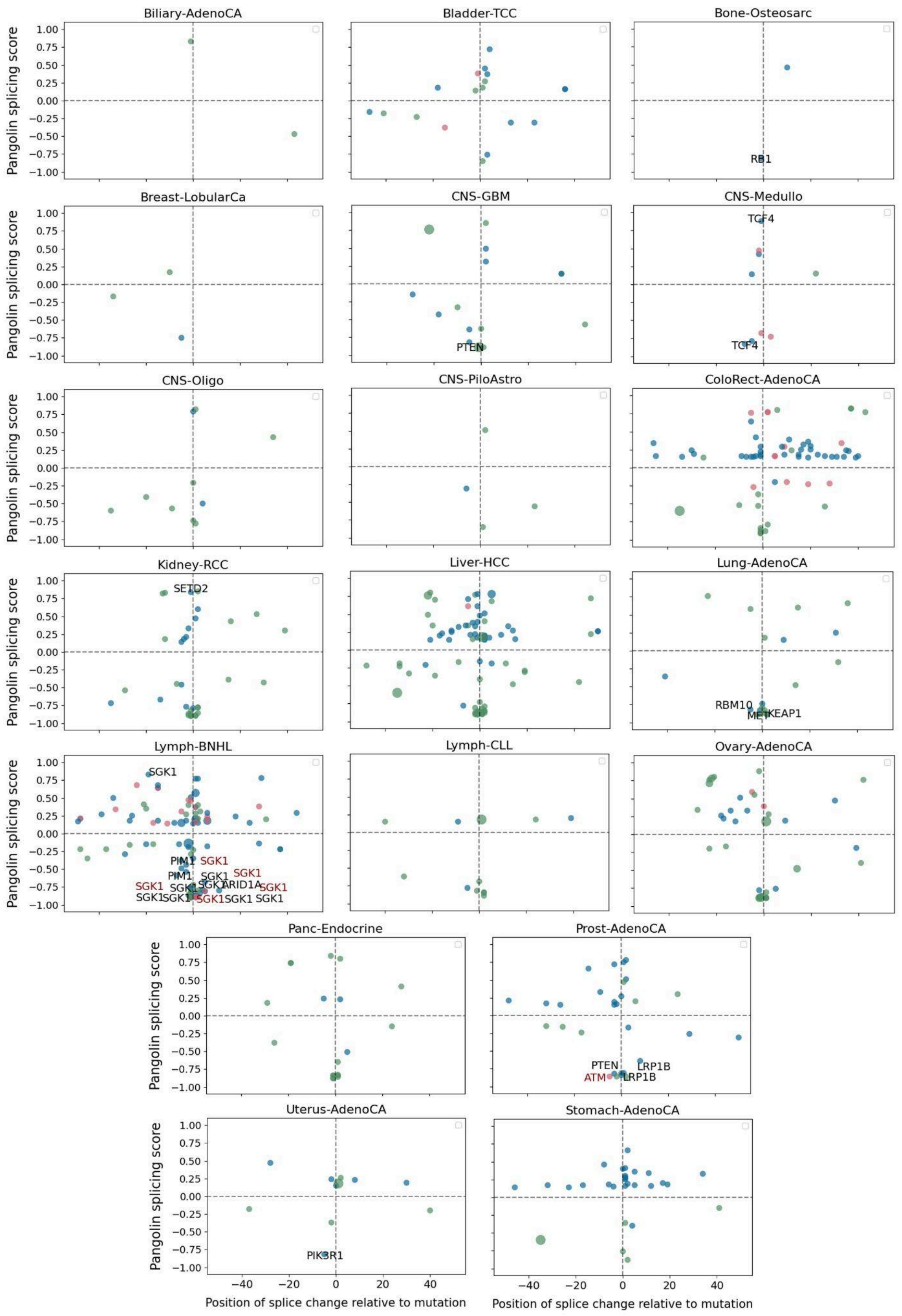


**Supplementary Figure 2: Distribution of splicing changes relative to the causative passenger and driver mutations. Genes harboring mutations causing large splicing changes (pangolin score >= 0.8) are annotated.**
